# Activation of group I mGluRs is required for heterosynaptic priming of long-term potentiation in mouse hippocampus

**DOI:** 10.1101/2024.09.27.615492

**Authors:** Laura A. Koek, Thomas M. Sanderson, John Georgiou, Graham L. Collingridge

**Author notes:** Department of Physiology and Biophysics, University of Colorado Denver, Aurora, CO, 80045, USA.

## Abstract

Metaplasticity involves changes in the state of neurons or synapses that influence their ability to generate synaptic plasticity. One form of heterosynaptic metaplasticity, known as synaptic tagging and capture (**STC**), has been intensively studied but the underlying mechanisms are not fully understood. In experiments using hippocampal slices prepared from C57BL/6J mice, we have examined the role of group I metabotropic glutamate receptors (**mGluRs**) in STC. We used a version of STC where a strong theta-burst stimulus (**TBS**), delivered to one set of Schaffer collateral-commissural pathway inputs to CA1, preceded a weak TBS delivered to a second independent set of inputs. We observed that, firstly, dual inhibition of mGluR1 and mGluR5, using YM 298198 and MTEP respectively, did not affect a form of protein synthesis-independent LTP (**LTP1**), but substantially inhibited a form of protein synthesis-dependent LTP (**LTP2**). Secondly, these inhibitors prevented the small heterosynaptic potentiation, which is often associated with LTP2. Thirdly, STC was abolished when these antagonists were applied either during the strong (priming) TBS or during the subsequent weak TBS at the independent pathway. It is proposed that the activation of group I mGluRs serves as a trigger for local protein synthesis both during the strong and weak TBS and, as such, are an integral part of the STC process. STC is involved in associative learning and memory, a cognitive function that is disrupted in many brain disorders including Alzheimer’s disease.

## INTRODUCION

Synaptic plasticity is extensively studied because it is a physiological process considered to be the central mechanism underlying learning and memory and its dysregulation leads to dementia and other brain disorders. The most extensively studied form of synaptic plasticity is long-term potentiation (**LTP**) at the excitatory synapses between CA3 and CA1 pyramidal neurons in the hippocampus (Bliss & Collingridge, 1993). However, despite the importance placed on understanding this process, there are still gaps in our knowledge. One such knowledge gap is in the relationship between homosynaptic and heterosynaptic plasticity, where plasticity is restricted to the activated synapses or spreads to neighbouring synapses, respectively.

LTP at CA3-CA1 synapses exists in both protein synthesis-independent and protein synthesis-dependent forms, sometimes referred to as LTP1 and LTP2, respectively (Bliss & Collingridge, 1993). LTP1 is readily induced by a single episode of high-frequency stimulation, such as a tetanus or theta-burst stimulation (**TBS**). LTP1 is also induced by multiple episodes of high-frequency stimulation, when the inter-episode interval (**IEI**) is on the order of tens of seconds. However, if the IEI is spaced out in the time frame of minutes, LTP2 is induced in addition to LTP1. The induction of both LTP1 and LTP2 requires the activation of NMDARs. However LTP2, but not LTP1, additionally requires the activation of calcium-permeable AMPARs (**CP-AMPARs**; Park et al., 2016, 2019; Park, Kang, et al., 2021), calcium-induced calcium release (**CICR**; Koek, Sanderson, et al., 2024), PKA (Frey et al., 1993; Matthies & Reymann, 1993), and *de novo* protein synthesis (Frey et al., 1988). The reason for two distinct forms of LTP at CA1 synapses is unknown, though one possibility is that, although LTP1 can last for at least a few hours (Bortolotto & Collingridge, 2000), LTP2 is the more persistent form of LTP that involves long-lasting structural changes at the synapse.

An interesting property of LTP is that its magnitude and pharmacological properties can be affected by prior activation of the synapses. In one such example, a brief tetanus altered pharmacological properties of LTP, via a process termed the molecular switch (Bortolotto et al., 1994). This priming effect did not require the activation of NMDARs but rather depended on the activation of mGluR5 (Bortolotto et al., 2005). In a closely related study it was shown that similar priming resulted in a larger LTP (Cohen & Abraham, 1996) due to a process coined metaplasticity (Abraham & Bear, 1996). This enhanced LTP, which was additive with the unprimed LTP, required *de novo* protein synthesis (Raymond et al., 2000). The unprimed potentiation can therefore be equated with LTP1 and the primed potentiation with LTP2, as described previously (Collingridge & Abraham, 2022). In these studies (Bortolotto et al., 1994; Raymond et al., 2000), the priming was homosynaptic, since an independent pathway was not altered in pharmacological sensitivity or magnitude.

The most extensively studied form of heterosynaptic metaplasticity is termed synaptic tagging and capture (**STC**). Here the priming of one input results in a greater LTP in an independent input (Frey & Morris, 1997; Sajikumar & Abel, 2024). Typically the priming is induced by a “strong” stimulus (defined as one that induces LTP2) and this enables a “weak” stimulus (defined as one that ordinarily induces only LTP1) at an independent pathway to additionally induce LTP2. In the canonical model, the stimulus at the strong input triggers widespread protein synthesis and the plasticity related proteins/products (**PRPs**) are captured by independent inputs that have been tagged by the ‘weak” stimulus (Redondo & Morris, 2011). In a modification to this model (Park, Kang, et al., 2021), we have proposed that the necessary *de novo* protein synthesis requires the synaptic activation of CP-AMPARs that are inserted into the synapse during the “strong” stimulus. We suggested that the synaptic tagging is achieved by the insertion of CP-AMPARs at perisynaptic sites at independent inputs. Thereafter, the “weak” stimulus drives these CP-AMPARs into the synapse to trigger local protein synthesis which generates LTP2, in addition to LTP1, at these inputs (see Koek, Park, et al., 2024).

Despite the established role of mGluRs in homosynaptic priming, it is not known whether the activation of mGluRs is also required for heterosynaptic priming. Therefore, in the present study we have addressed whether mGluRs are involved in this form of metaplasticity. Here we report that group I mGluRs are involved in LTP2, heterosynaptic potentiation and heterosynaptic priming of LTP. Group I mGluRs are therefore part of a functional synaptic metaplasticity module that also involves NMDARs, CP-AMPARs, CICR, PKA and *de novo* protein synthesis (Koek, Park, et al., 2024).

## RESULTS

To explore the effects of antagonism of group I mGluRs on LTP we simultaneously inhibited mGluR5 and mGluR1 using MTEP (1 µM) and YM 298198 (**YM**; 1 µM), respectively, and tested three different LTP induction protocols. Antagonism of both group I mGluRs, did not significantly affect the induction of LTP induced by either a weak TBS (wTBS; fig. 1A-B) or compressed TBS (cTBS; fig. 1C-D). Quantified 60 min following a wTBS, the level of LTP was 27± 6 % (n = 6) and 22 ± 9% (n = 6) above baseline (100 %) for control slices and those treated with MTEP+YM, respectively (p = 0.676). Quantified 120 min following a cTBS, the level of LTP was 41± 6 % (n = 10) and 44 ± 4% (n = 6), respectively (p = 0.781).

**Figure 1.**
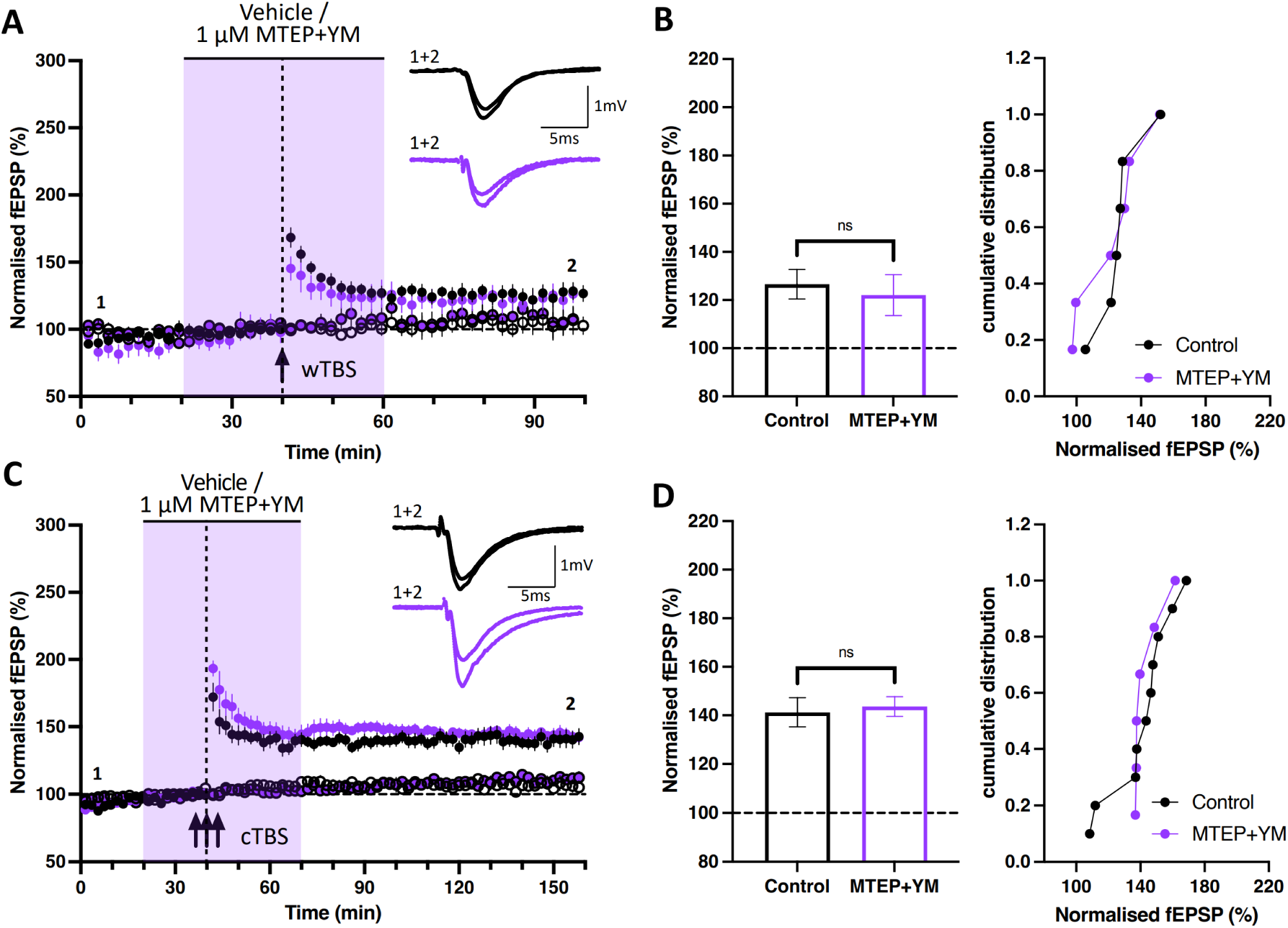
Group 1 mGluRs are not required for LTP1. **(A)** Time-course plot of CA3-CA1 fEPSP synaptic responses showing that 1 µM MTEP+YM 298198 (YM) treatment (purple symbols) did not significantly affect LTP induced by a weak TBS (wTBS) at input S0 (filled circles). The control independent input (S1, open circles) was also unaffected. **(B)** The mean (± SEM) level of LTP in S0, with (right) cumulative probability plot of the LTP obtained from each individual experiment. **(C-D)** The equivalent experiments for cTBS, compressed TBS. In these and subsequent figures, data from controls are black or white open circles (S0 and S1, respectively) and data from treated slices are colour-coded. The duration of compound application is denoted by the shaded rectangle. Sample fEPSP synaptic responses (to the upper right of the time-course plots) are the average of 4 consecutive responses obtained from a representative experiment at the times indicated by 1 and 2. * p<0.05, **p<0.01, ***p<0.001 (see text for actual p values).

However, the application of MTEP+YM significantly reduced the level of LTP when these antagonists were present during the sTBS (fig. 2). Quantified 120 min following the delivery of sTBS to input S0, the level of LTP was 73 ± 13 % (n = 6; fig. 2A, upper) and 23 ± 3 % (n = 6; fig. 2B, upper), respectively (p = 0.002). In contrast when YM+MTEP were applied starting 10 min after completion of the sTBS protocol, the level of LTP of 45 ± 9 % (n = 4; fig. 2C, upper) was not significantly different from control (p = 0.089). The sTBS results for input S0 are summarised as histograms and as cumulative probability plots, which show the individual data points for each experiment (fig. 2D).

**Figure 2.**
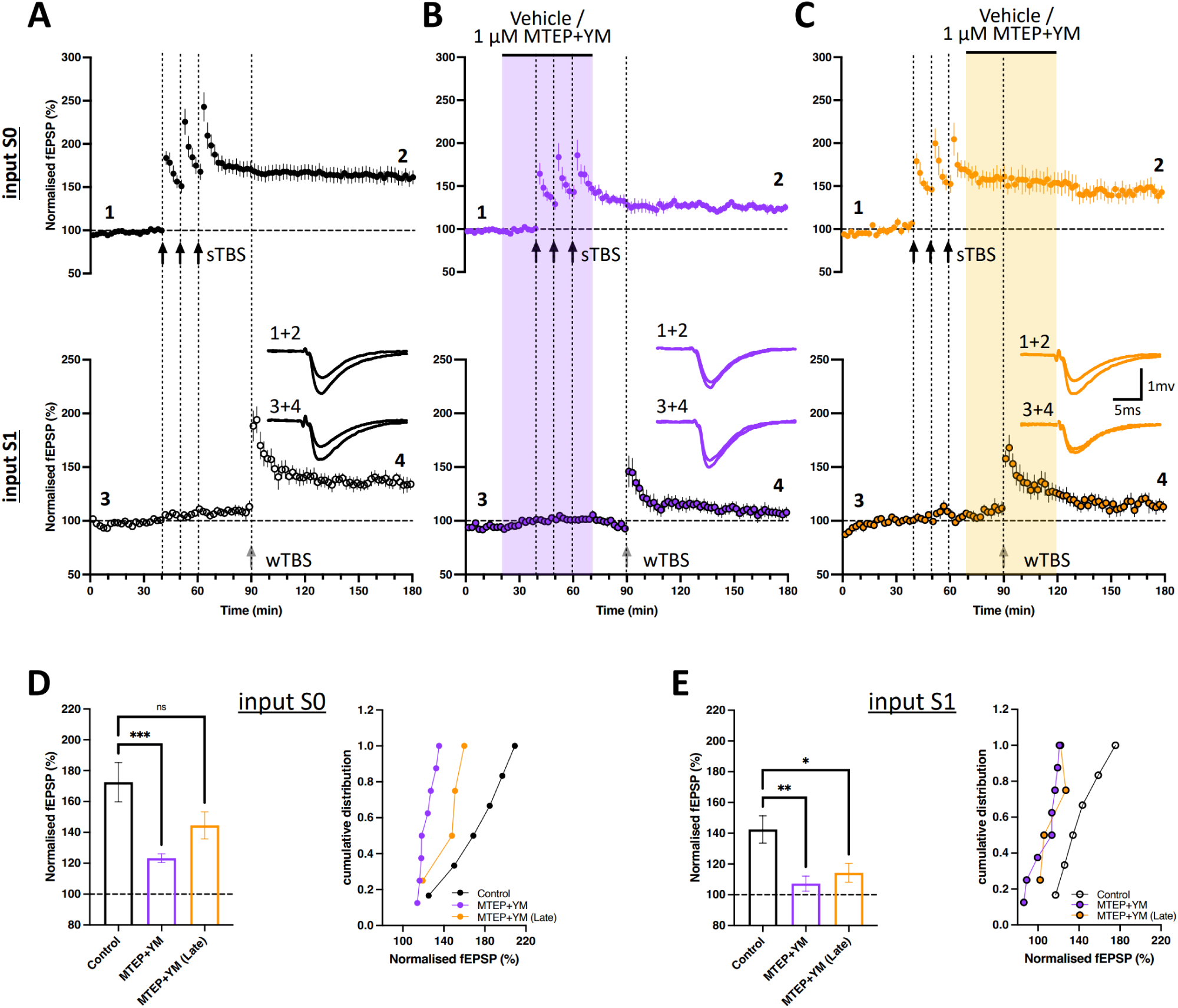
Role of Group I mGluRs in LTP2. **(A)** STC control experiments where sTBS at input S0 (upper time-course data) precedes the wTBS at input S1 (lower data) by 30 min (i.e. primed). **(B)** When 1 µM MTEP+YM was applied for a 50 min period (purple column) that overlapped the sTBS delivery, these mGluR5/1 inhibitors greatly reduced homosynaptic LTP on input S0 and STC on independent input S1. **(C)** Time-course plot showing that a late application of 1 µM MTEP+YM after the sTBS only inhibited STC on input S1 significantly. **(D)** Corresponding LTP (left) and cumulative distribution plots (right) at t = 170-180 min for input S0. **(E)** Corresponding LTP and cumulative probability plots at t = 170-180 min for input S1.

In terms of heterosynaptic effects, we quantified the level of total potentiation (relative to the initial baseline) 90 min following a wTBS at input S1, which was delivered 30 min following the sTBS at input S0. Compared to the control level of potentiation (42 ± 9 %; n = 6; fig. 2A, lower), LTP at input S1 was significantly inhibited when MTEP+YM were applied either during the sTBS at input S0 (7 ± 5 %; n = 8; p = 0.003; fig. 2B) or during the wTBS at input S1 (14 ± 6 %; n = 4; p = 0.035; fig. 2C, lower). The wTBS results for input S1 are summarised as histograms and as cumulative probability plots in fig 2E.

To determine the relative roles of mGluR1 and mGluR5 we compared the effects of YM versus MTEP in interleaved experiments, which also included a new set of controls. Compared to the control level of LTP at input S0 (48 ± 10 %; n = 7; fig. 3A, upper) the effects of neither MTEP (62 ± 11 %; n = 6; p = 0.552; fig. 3B, upper) nor YM (79 ± 12 %; n = 5; p = 0.111; fig. 3C, upper) were significantly different. Similarly, for the heterosynaptic effect at input S1, the levels of LTP for control of 52 ± 7 % (n = 7; fig. 3A, lower), MTEP-treated (30 ±8 %; n = 6; p=0.117; fig. 3B, lower) and YM-treated (38 ± 10 %; n = 5; p=0.395; fig. 3C, lower) slices were not statistically different.

**Figure 3.**
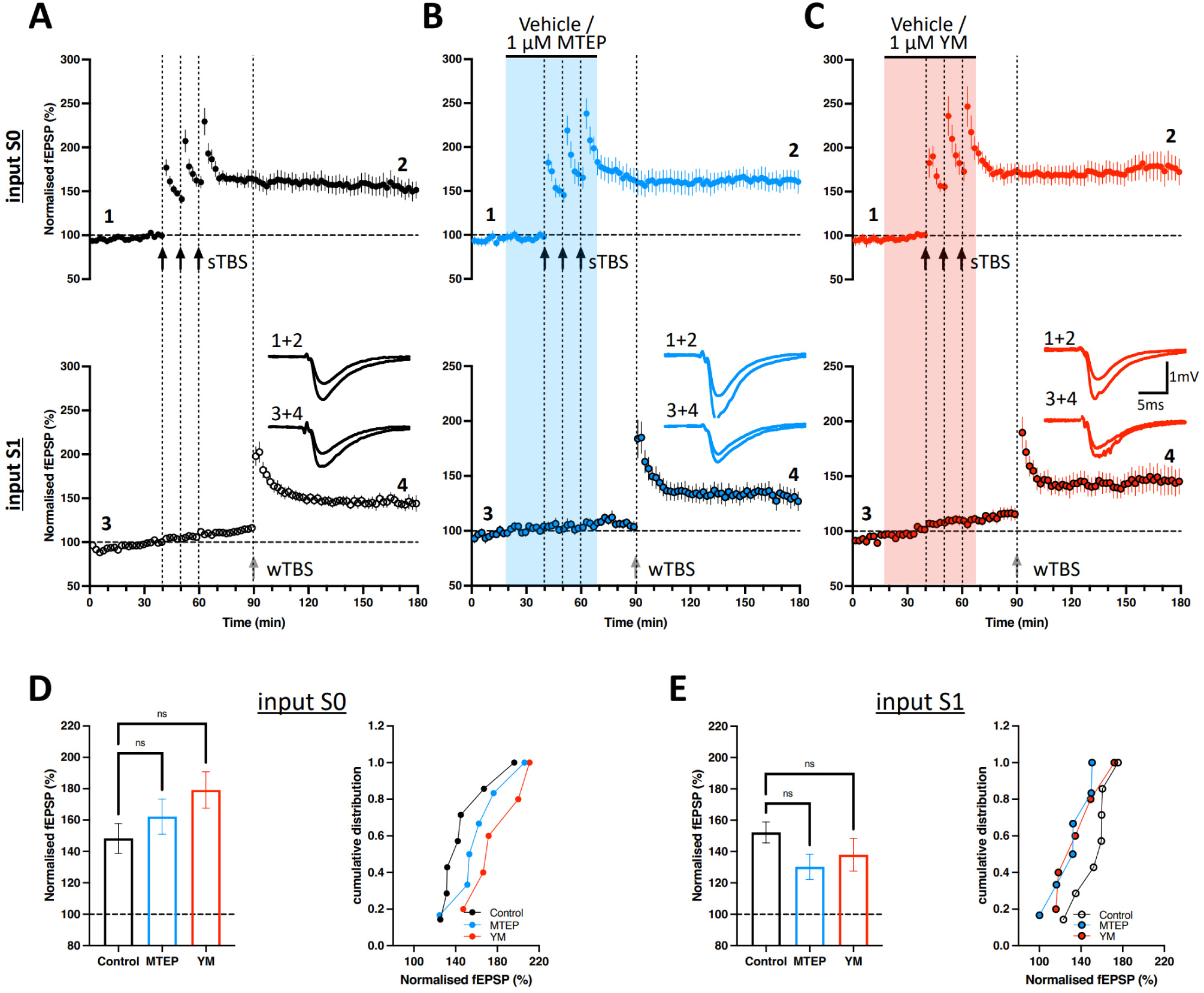
Differential roles of mGluR1 and mGluR5 in LTP2. **(A)** Interleaved control experiments where sTBS at input S0 (upper time-course data) preceded the wTBS at input S1 (lower data) by 30 min (i.e. primed). **(B)** Application of 1 µM MTEP (to block mGluR5) during the sTBS did not affect homosynaptic LTP on input S0 or heterosynaptic potentiation on input S1. **(C)** When 1 µM YM (to block mGluR1) was applied during the sTBS, this treatment did not affect homosynaptic LTP on input S0 or heterosynaptic potentiation on input S1. **(D)** Plots of the corresponding LTP (left) and cumulative distribution (right) at 170-180 min for input S0 according to treatment. **(E)** Corresponding LTP and cumulative probability plots at 170-180 min for input S1.

The total synaptic potentiation at S1 comprises three mechanistically distinct components, (i) the underlying LTP that would be observed in the absence of priming (i.e., LTP1), (ii) additional LTP due to heterosynaptic priming (i.e., to generate an additional LTP2 rather than only LTP1) and also (iii) a modest heterosynaptic potentiation (i.e. following sTBS at S0 but in the absence of any TBS at S1). This heterosynaptic potentiation was observed in some but not all the conditions. For example, in experiments where the effects of MTEP+YM applied together were addressed (fig. 4A-C, G), significant heterosynaptic potentiation was observed in interleaved controls (18 ± 8.2 %; n = 6; p=0.039, paired t-test between t = 30-40 min and t = 80-90 min data, here and in the following comparisons; fig. 4A), as well as when MTEP+YM were co-applied following the sTBS (14 ± 6 %; n = 4; p = 0.016; fig. 4C), but not when MTEP+YM were co-applied during the sTBS (−7 ± 3; n = 8; p = 0.430; fig. 4B). When the effects of the group I mGluR antagonists applied individually were addressed (fig. 4D-F, H), heterosynaptic potentiation was observed in interleaved controls (14 ± 2%; n = 7; p = 0.009; fig. 4D), when YM was applied alone (18 ± 6 %; n = 5; p = 0.041; fig. 4F, H), but not when MTEP was applied alone (3 ± 4 %; n = 6; p = 0.957; fig. 4E, H).

**Figure 4.**
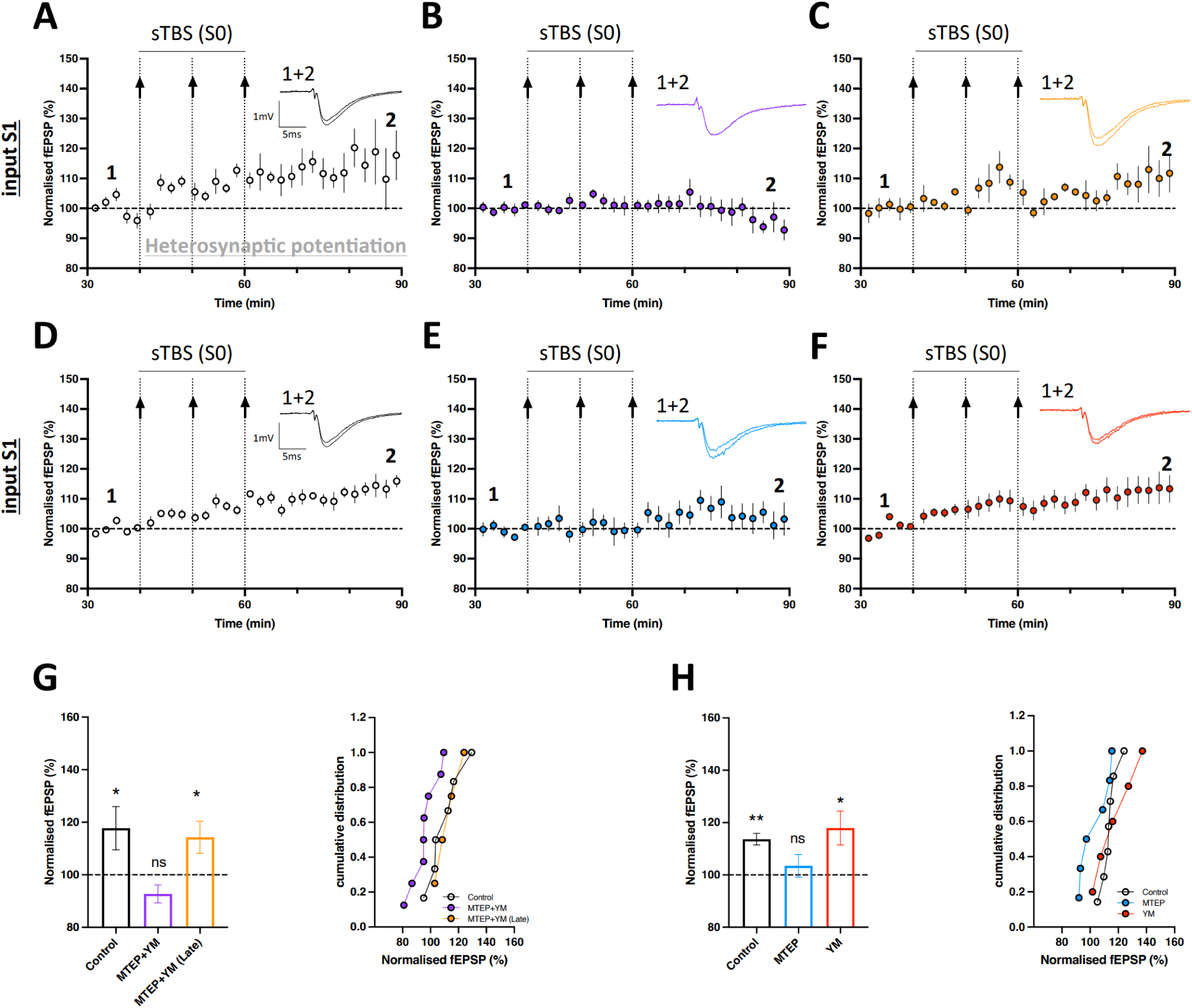
The role of mGluR1 and mGluR5 in heterosynaptic potentiation. Synaptic response time-course data from fig. 2A-C <u>input S1</u> were renormalised to t = 30-40 min, to get a true estimate of heterosynaptic potentiation: **(A)** control group, **(B)** effect of 1 µM MTEP+YM treatment, and (C) late application of 1 µM MTEP+YM. In the next row of data (D-F), the corresponding experimental input S1 data from fig. 3A-C were similarly renormalised to t = 30-40 min baseline: (D) Control, **(E)** 1 µM MTEP, and **(F)** 1 µM YM. **(G)** Bar plots for the corresponding level of potentiation (left) at t = 80-90 min for data in A-C with corresponding cumulative distribution plots (right)**. (H)** Similar analysis of the LTP (left) for D-F data, with the corresponding cumulative probability distribution (right).

To quantify the level of LTP at input S1 following heterosynaptic priming (at input S0) we renormalised the data to the baseline over the 10 min immediately preceding the wTBS (at input S1). By this time, heterosynaptic potentiation had stabilised and so its subtraction enables a good estimate of LTP1 + LTP2. Compared to the control level of LTP at input S1 (26 ± 5 %; n = 6; fig. 5A), there was significant inhibition by MTEP+YM when applied both during ( 9 ± 4 %; n = 8; p = 0.021; fig. 5B) and following (4 ± 3 %; n = 4; p = 0.011; fig. 5C) the delivery of sTBS to input S0. In contrast, compared to the level of LTP at input S1 (34 ± 6 %; n = 7; fig. 5D), in another set of control data, there was no significant effect of either MTEP ( 20 ± 5 %; n = 6; p = 0.189; fig. 5E) or YM (29 ± 5 %; n = 5; p = 0.774; fig. 5F). These collective data are graphed in fig 5G-H.

**Figure 5.**
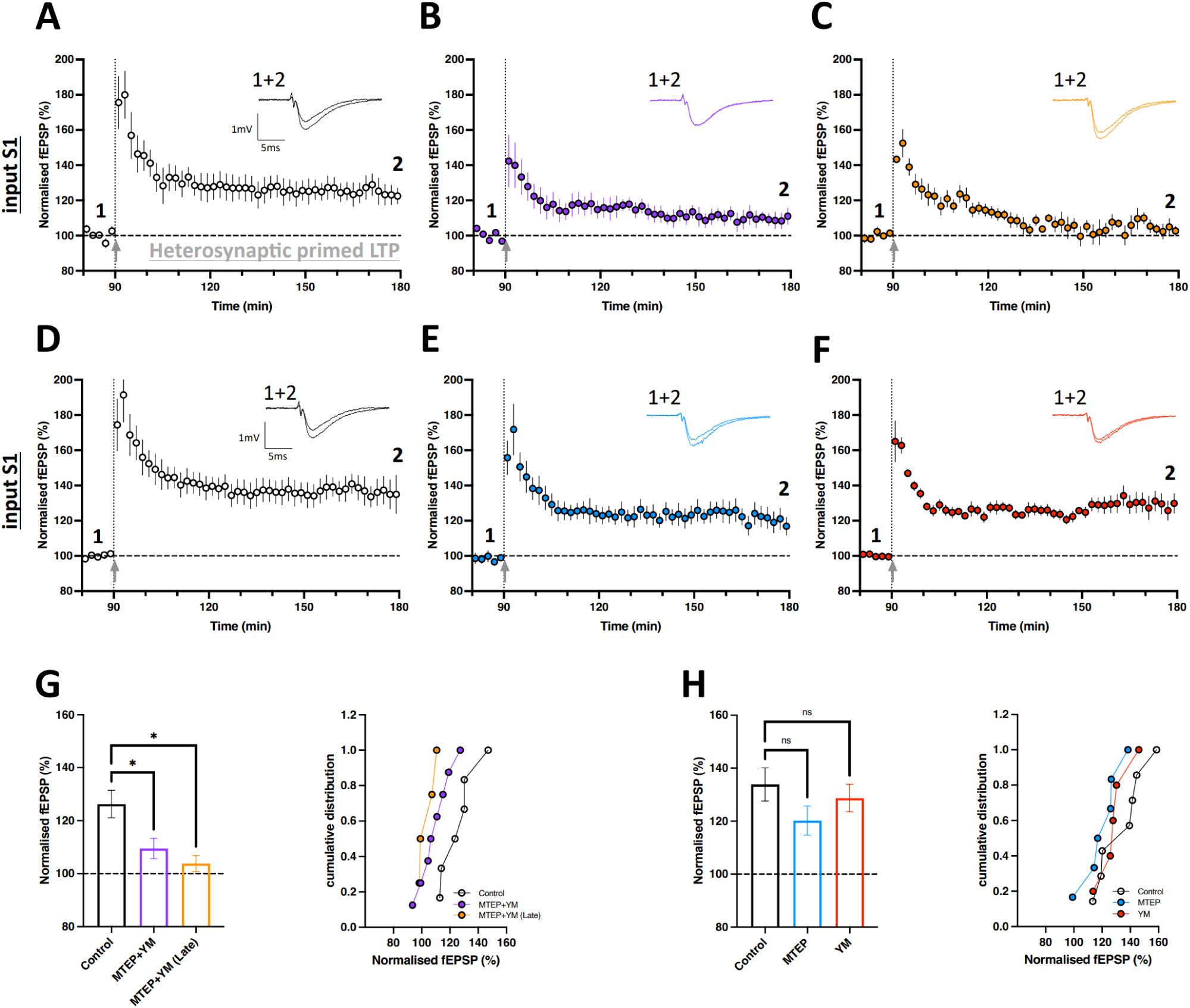
The role of mGluR1 and mGluR5 in heterosynaptic primed LTP. Synaptic response time-course data from fig. 2A-C <u>input S1</u> were renormalised to t = 80-90 min, to get a true estimate of heterosynaptic primed LTP: **(A)** control group, **(B)** effect of 1 µM MTEP+YM treatment, and (C) late application of 1 µM MTEP+YM. In the next row of data (D-F), the corresponding experimental input S1 data from fig. 3A-C were similarly renormalized to t = 30-40 min baseline: (D) Control, **(E)** 1 µM MTEP, and **(F)** 1 µM YM. **(G)** Bar plots for the corresponding level of potentiation (left) at t = 170-80 min for data in A-C with corresponding cumulative distribution plots (right). **(H)** Similar analysis of the LTP (left) for D-F data, and the corresponding cumulative distribution plots (right).

## DISCUSSION

The principal finding of the present study is that group I mGluRs are important for a variety of forms of synaptic plasticity and metaplasticity in the hippocampus. A schematic of their roles is shown in fig. 6. We have observed inhibition of LTP2, heterosynaptic potentiation and heterosynaptic priming of LTP1 but no effect on unprimed LTP1. LTP1 and LTP2 are distinguished by their independence and dependence, respectively, on *de novo* protein synthesis, PKA, CP-AMPARs and CICR (Y. Huang & Kandel, 1994; Koek, Sanderson, et al., 2024; Park et al., 2013, 2016). Heterosynaptic potentiation (Park et al., 2019) and heterosynaptic priming of LTP1 (Koek, Park, et al., 2024) also require the activation of these same signalling cascades, which can collectively be considered an LTP2 synaptic plasticity module. It can therefore be concluded that group I mGluRs are associated with this same module.

**Figure 6.**
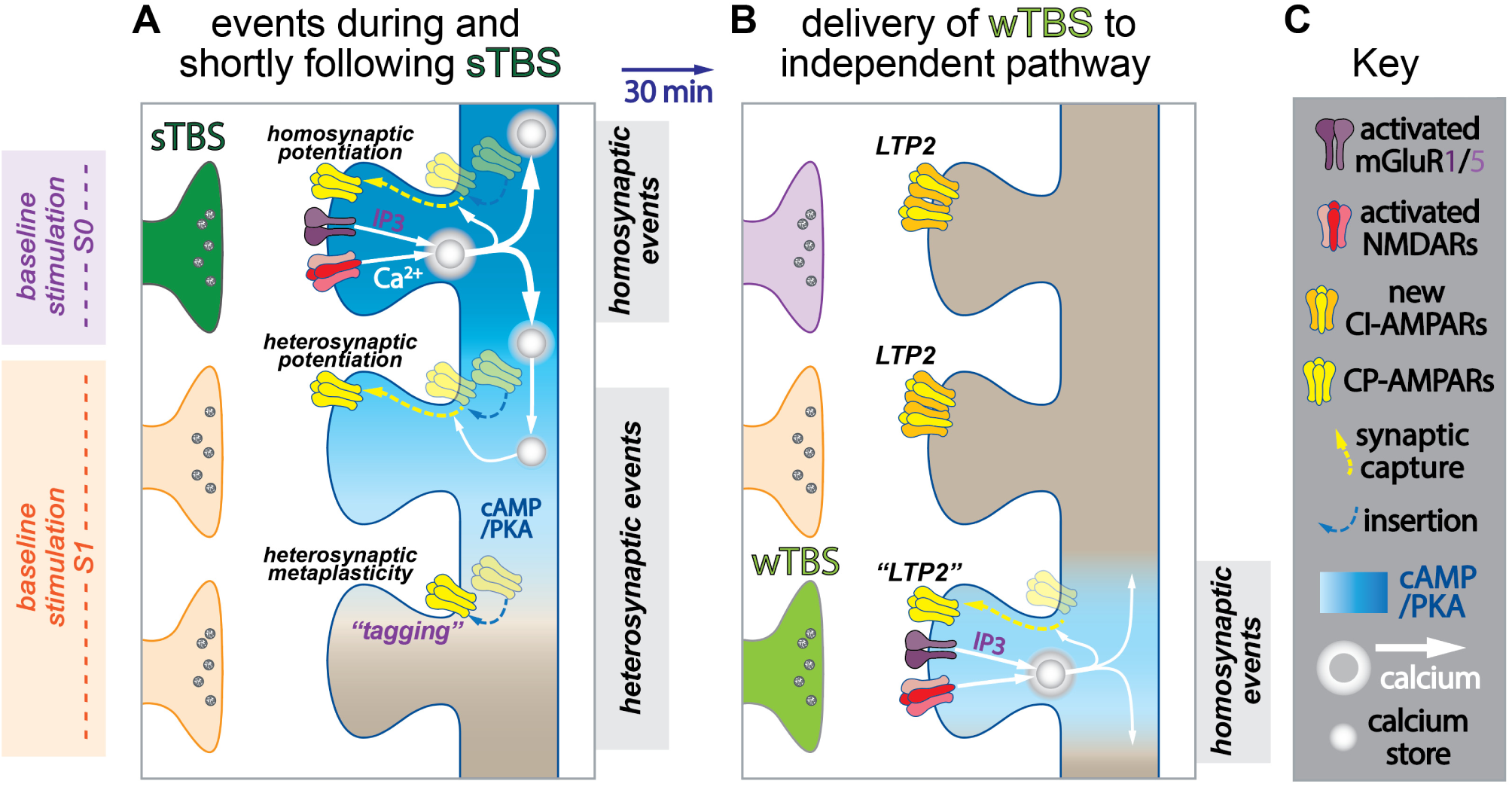
A potential scheme to explain LTP2, heterosynaptic potentiation and STC based on differential spread of Ca^2+^ and cAMP. **(A)** Three spines are depicted, of which one is activated by stimulation of input S0 (upper row) and another by stimulation of input S1 (third row). These represent a much larger population of intermingled spines on a given dendritic segment. Delivery of a sTBS (“strong stimulus”) to input S0 results in homosynaptic LTP1 (not illustrated) and homosynaptic LTP2. The latter is due to the transient synaptic insertion of calcium-permeable AMPARs (CP-AMPARs) driven by a two-step process involving cAMP/PKA (blue gradient) and Ca^2+^ (white arrows). The Ca^2+^ signal derives from calcium-induced Ca^2+^ release (CICR) which is stimulated by the Ca^2+^ entering via NMDARs and the IP3 generated from group I mGluR activation. Subsequent baseline stimulation drives Ca^2+^ through these receptors to trigger the exchange of the CP-AMPARs for a greater number of calcium-impermeable AMPARs (CI-AMPARs). Critical to heterosynaptic events is the activation of PKA in surrounding synapses, most probably due to the diffusion of cAMP that drives CP-AMPARs into the plasma membrane. In some instances, these are driven into the synapse to generate heterosynaptic potentiation, and in others they remain at perisynaptic sites to tag synapses. **(B)** At heterosynaptic sites where CP-AMPARs are synaptically inserted (see the spine in the second row from panel A), subsequent baseline stimulation drives Ca^2+^ through these receptors to trigger their replacement with a greater number of CI-AMPARs, resulting in a stable heterosynaptic LTP2. At tagged synapses (i.e. where CP-AMPARs are perisynaptically located, see the spine in the third row of panel A), a subsequent wTBS (“weak stimulus”) is able to drive them into the synapse to trigger LTP2 in exactly the same manner as described in panel A. This process again requires CICR and hence activation of group I mGluRs. In addition to aiding the spread of Ca^2+^ via CICR, group I mGluRs also help trigger *de novo* protein sythesis. Note that these functional chnages are accompanied with structural changes, which are not shown for simplicity. **(C)** Key. Note that for simplicity in A and B, AMPAR clusters involved in basal synaptic responses and LTP1 are not illustrated. Only NMDAR clusters and mGluRs activated during the TBS are illustrated (Modified from Koek, Sanderson, et al., 2024).

### Role of mGluRs in LTP

Many studies have investigated the role of mGluRs in LTP in a variety of pathways in the brain (Anwyl, 2009). Here we restrict our discussion to CA3-CA1 synapses, where most information has been gathered, yet the situation is not straightforward. Notably, two forms of LTP that can co-exist at the same population of synapses were defined by their sensitivity and insensitivity to MCPG (Bashir et al., 1993; Bortolotto et al., 1994), an antagonist that acts on some, but not all, mGluR subtypes. The subsequent development of mGluR1 and mGluR5 selective antagonists has enabled their individual roles to be explored. With respect to LTP1 (i.e., protein synthesis-independent LTP) there have been mixed conclusions. For example, LTP in a constitutive mGluR1 KO was reported to be reduced (Aiba et al., 1994) or unaffected (Conquet et al., 1994). In a constitutive mGluR5 KO, LTP was reported to be partially inhibited (Bortolotto et al., 2005; Lu et al., 1997). In pharmacological experiments using mGluR1 selective antagonists, LTP was inhibited (Foster et al., 2018; Francesconi & Duvoisin, 2004; Neyman & Manahan-Vaughan, 2008; Tigaret et al., 2016, 2018) or unaffected (Latif-Hernandez et al., 2016). Using mGluR5 selective antagonists LTP was inhibited (Francesconi & Duvoisin, 2004; Hagena et al., 2022; Kwag & Paulsen, 2012; Latif-Hernandez et al., 2016; Neyman & Manahan-Vaughan, 2008) or unaffected (Bortolotto et al., 2005; Doherty et al., 2000; Tigaret et al., 2016). In the present study the combined application of selective mGluR1 and mGluR5 antagonists also had no significant effect on LTP1, induced by two different protocols.

Regarding LTP2, the conclusions have also been mixed. In one study, either mGluR1 or mGluR5 antagonists reduced LTP (Fan, 2013) and in another, inhibition of mGluR5 but not mGluR1 was effective (Francesconi & Duvoisin, 2004). Related to the latter study, a mGluR5 positive allosteric modulator (PAM) selectively enhanced LTP2 (Kroker et al., 2011). In the present study, the combined application of selective mGluR1 and mGluR5 antagonists reduced LTP when present during sTBS induction. Since MTEP+YM did not affect LTP1, that was induced with either a wTBS or cTBS, the residual LTP that was unaffected by MTEP+YM treatment can be attributed to LTP1, which is invariably induced along with LTP2 with this sTBS protocol. It seems likely therefore that the combined treatment with MTEP+YM eliminated LTP2. When applied following the sTBS, MTEP+YM had no significant effect on LTP. These data suggest that activation of group I mGluRs are required for the induction, but not the maintenance or expression of LTP2. In contrast, to the effectiveness of combined inhibition of mGluR1 and mGluR5, neither MTEP nor YM applied alone had any effect on LTP2. This could be because either mGluR1 and mGluR5 are both able to induce LTP2, such that inhibition of just one subtype is ineffective, or because the group I mGluR is a mGluR1/mGluR5 dimer and can be activated by L-glutamate binding to one or other of the subunits, for which precedence exists (McCullock & Kammermeier, 2021; Ng et al., 2023; Belkacemi et al., 2024).

Although there has been considerable disagreement regarding the roles of group I mGluRs in LTP at these synapses, a few firm conclusions can be drawn: (i) activation of group I mGluRs is not essential for the induction of LTP. This conclusion is supported by the observations that LTP can be readily induced following pharmacological inhibition of all mGluR subtypes, using LY341495 (Fitzjohn et al., 1998), and that LTP can be induced in an mGluR1/mGluR5 double constitutive KO (Bortolotto et al., 2005). (ii) both mGluR1 and mGluR5 can strongly regulate the induction of LTP. Why the variability? With respect to LTP1, one possibility is that it depends on the level of depolarisation generated by the synaptic inputs. A critical level is necessary to reduce the Mg^2+^ block of NMDARs and this might require mGluRs to facilitate the synaptic activation of NMDARs under some circumstances but not others (Bliss & Collingridge, 1993). This, in turn, might relate to the number of afferents activated and/or the induction protocol used. Also, conditions, such as a single or compressed induction protocol, that would be predicted to induce LTP1 alone could also induce LTP2 due to neuromodulation. Indeed, this would be expected under conditions of elevated PKA activity, where a single episode of TBS readily induces LTP2 (e.g., Park, Georgiou, et al., 2021). Examples of this include level of modulation by dopamine (Huang & Kandel, 1995) and the level of stress the animal experiences prior to slice preparation (Whitehead et al., 2013).

Our tentative conclusion is that an mGluR1 homodimer facilitates, but is not essential for, the induction of LTP1 and that an mGluR1/mGluR5 heterodimer is required for LTP2. This latter conclusion is consistent with the established roles of group I mGluRs, to trigger local *de novo* protein synthesis (Raymond et al, 2000).

### Role of mGluRs in homosynaptic priming

The initial controversy surrounding the ability of MCPG to inhibit the induction of LTP, with a few reports reporting no effect (Chinestra et al., 1993; Selig et al., 1995), led to the discovery that activation of mGluRs could prime synapses to alter their subsequent sensitivity to mGluR antagonism (Bortolotto et al., 1994). In another manifestation of this phenomenon, mGluR priming leads to enhanced LTP in the absence of mGluR antagonism (Cohen & Abraham, 1996). Since only the primed LTP is sensitive to inhibitors of protein synthesis (Raymond et al., 2000), it can be concluded that the unprimed LTP is LTP1 and the primed LTP is LTP2, as discussed previously (Collingridge & Abraham, 2022). This priming phenomenon, which is restricted to activated synapses (Bortolotto et al., 1994; Cohen & Abraham, 1996), gave rise to the term of metaplasticity (Abraham & Bear, 1996). Several properties of homosynaptic priming have been identified. The priming does not require activation of NMDARs (Bortolotto et al., 1994; Cohen & Abraham, 1996), mGluR1 or mGluR7 but is totally dependent on activation of mGluR5 (Bortolotto et al., 2005), for which only a few stimuli are required (Bortolotto et al., 2008). It involves activation of PLC (Cohen & Abraham, 1996), CaMKII (Bortolotto & Collingridge, 1998) and PKC, but not PKA (Bortolotto & Collingridge, 2000).

### Synaptic Tagging and Capture

One form of heterosynaptic metaplasticity, termed synaptic tagging and capture (STC), has been extensively studied (Sajikumar & Abel, 2024). In the original description of STC it was shown that a strong stimulus (defined as one that induced protein synthesis-dependent LTP; i.e., LTP2) to one input induced augmentation of the LTP induced by a weak stimulus (defined as one that induced protein synthesis-independent LTP; i.e., LTP1) to an independent input (Frey & Morris, 1997). In addition, the strong stimulus could induce a small heterosynaptic potentiation. In the present investigation, we have used a similar protocol and observed both heterosynaptic potentiation and heterosynaptic metaplasticity. Our “strong” stimulus comprised 75 pulses delivered as a spaced TBS (sTBS) protocol that induces LTP1 plus LTP2, and our “weak” stimulus (wTBS) comprised of 25 pulses that induced LTP1 alone. Since the “strong” sTBS preceded the wTBS, it is a form of heterosynaptic priming. In the present study we therefore use the following terminology: the total heterosynaptic effect induced by the “strong stimulus” that is commonly referred to as STC is composed of two components, heterosynaptic potentiation and heterosynaptic priming. These are then additive to the LTP1 that the wTBS evokes.

### Role of mGluRs in heterosynaptic potentiation

In the present study we found that heterosynaptic potentiation was readily induced. However when group I mGluR inhibitors were applied during, but not following the sTBS, it was not (fig. 4). For the inhibition of heterosynaptic potentiation, selective inhibition of mGluR5 was sufficient. The simplest explanation is that during sTBS, mGluRs trigger LTP2 that is not restricted to homosynaptic inputs. Consistent with this notion that heterosynaptic potentiation is a spatial dendritic expansion of LTP2 are the findings that it also requires activation of CP-AMPARs and *de novo* protein synthesis (Park et al., 2019). Two problems that arise from these observations is where are the group I mGluRs that mediate heterosynaptic potentiation located and how are they activated? One possibility is that they are activated at the synapses that receive the sTBS and help trigger a signal that propagates to the heterosynaptic input; for example, they help trigger the CICR that is also required for heterosynaptic effects (Koek, Sanderson, et al., 2024). Their ability to generate inositol trisphosphate (IP3) is consistent with this idea. A second possibility is that they are activated at the heterosynaptic input by “spill-over” from the input receiving sTBS. A third possibility is that they are activated locally by the test stimuli. The last two possibilities may be facilitated by the withdrawal of perisynaptic astroglia that occurs in LTP (Hennenberger et al., 2020).

### Role of mGluRs in heterosynaptic priming

Inhibition of group I mGluRs abolished the effects of heterosynaptic priming, when applied during the wTBS. This result suggests that activation of mGluRs is required for both the priming of and induction of LTP2 in response to the weak induction protocol. The simplest explanation is that mGluRs are required to be activated by the sTBS to help prime the heterosynapse, potentially by enabling the rapid synthesis and/or perisynaptic insertion of CP-AMPARs, and are required again during the wTBS to help trigger local protein synthesis of primed inputs.

It is worth noting that we have used the terms “strong” and “weak” throughout to denote (i) the priming stimulus and (ii) the wTBS stimulus at the primed input, respectively. In reality we believe that it’s the timing of the stimuli rather than the strength *per se* that determines whether priming occurs or not. We found that 75 stimuli delivered as a sTBS induced priming whereas 75 identical stimuli delivered as a cTBS did not (Park et al., 2019). This priming correlates with the ability of the spaced, but not the compressed, protocols to lead to the synaptic insertion of CP-AMPARs, activation of PKA and triggering of *de novo* protein synthesis.

### Limitations of the study

The principal conclusion of the present study is that group I mGluRs are required for heterosynaptic metaplasticity, a principal component of the STC process. However, our study has only focused on the first 90 min of the primed LTP; further studies are required to investigate whether the mGluR-dependent priming effect extends to longer periods. Our study has also been restricted to a single priming protocol, where the strong stimulus precedes the weak stimulus. Again, additional experiments are required to investigate the STC process when the weak stimulus precedes the strong one (Frey & Morris, 1998).

### Concluding Remarks

Studies of homosynaptic metaplasticity and heterosynaptic metaplasticity have largely been conducted in isolation from one another. However, there are many similarities between these two forms of metaplasticity. Principally, the two forms of metaplasticity differ in whether they are homosynaptic or heterosynaptic in nature. As such we posit that heterosynaptic priming is essentially the same mechanism as homosynaptic priming but with the added requirement for a mechanism to overcome synapse specificity. We have proposed that the primary signal that extends plasticity beyond the activated synapses is cAMP, which activates PKA at neighbouring synapses (Tang & Yasuda, 2017). PKA then phosphorylates GluA1 to drive CP-AMPARs into the perisynaptic plasma membrane (see Koek, Park, et al., 2024). A subsequent wTBS, at an independent input, is able to drive the CP-AMPARs into the synapse where their activation by test pulse stimuli leads to *de novo* protein synthesis resulting in LTP2 (in addition to LTP1). This model, which builds upon the concept of clustered plasticity (Govindarajan et al., 2011), does not exclude other factors involved in heterosynaptic cross-talk, such as RhoA and Rac1 (Hedrick et al., 2016). It does, however, posit CP-AMPARs as the synaptic tag and mGluR5 as a principal trigger for PRP synthesis. As these receptors have been implicated in brain disorders that include Alzheimer’s disease, the findings herein are of relevance to identification of synaptic plasticity and STC mechanisms in health and disease.

## METHODS

### Slice preparation

Experiments were performed as described in Koek et al., 2024. Briefly, C57BL/6J male mice (10-12 weeks of age) were anaesthetized with isoflurane and euthanised by decapitation in accordance with the Canadian Council on Animal Care (CCAC) guidelines and an AUP approved by The Centre for Phenogenomics (Toronto, Canada) Animal Care Committee. Transverse hippocampal slices (400 µm) were prepared using a vibratome (VT1200S; Leica Biosystems, Canada). The cutting solution for the dissection contained (in mM) 124 NaCl, 3 KCl, 26 NaHC0_3_, 1.25 NaH_2_PO_4_, 10 D-glucose, 5 MgSO_4_ and 1 CaCl_2_. The CA3 region was cut with a scalpel blade to reduce upstream neuronal excitability and slices were transferred to an incubation chamber containing recording solution (artificial cerebrospinal fluid, ACSF (in mM) 124 NaCl, 3 KCl, 26 NaHCO_3_, 1.25 NaH_2_PO_4_, 2 MgSO_4_, 10 D-glucose and 2 CaCl_2_ (bubbled with 95% O_2_ and 5% CO_2_). Slices were allowed to recover at 32 °C for 30 min after sectioning and for a minimum of 1 h at 22 °C before recordings were obtained.

### Electrophysiology

Hippocampal slices were perfused at 2 mL/min with the oxygenated ACSF at 30 °C. Two bipolar electrodes were positioned in the stratum radiatum on either side and equidistant to a recording electrode. Two independent SCC pathways (denoted S0 and S1) were stimulated alternately using a constant current stimulator (0.033 Hz frequency, pulse width 0.1 ms; STG 4002, Multichannel systems, Germany) to evoke synaptic responses. Signals were amplified with a Multiclamp 700B (Molecular Devices, USA) and digitized with a BNC-2110 (National Instruments, Canada) A/D board, at a sampling rate of 20 kHz. The independence of the two inputs was verified by the absence of heterosynaptic paired-pulsed facilitation, using an inter-pulse interval of 50 ms. Throughout the experiments, paired pulses were applied within each pathway (50 ms inter-stimulus interval). The initial slope of the evoked field excitatory postsynaptic potential (fEPSP) (V/s) was monitored and analysed using WinLTP 11 (WinLTP Ltd, UK) (Anderson & Collingridge, 2007).

Following a stable baseline of at least 20 min, LTP was induced using theta-burst stimulation (TBS) delivered at the same current intensity and pulse width. An episode of TBS comprised 5 bursts at 5 Hz with each burst composed of 5 pulses at 100 Hz (25 pulses in total). To induce LTP 1, either a weak (w) TBS or compressed (c) TBS was used where wTBS comprised one episode of TBS (25 pulses) and cTBS comprised three TBS episodes, delivered with an inter-episode interval (IEI) of 10 s (75 pulses in total). To induce LTP2, a spaced (s) TBS comprising three TBS episodes, with an IEI of 10 min (75 pulses total) was used. The protocol for studying heterosynaptic metaplasticity was similar to that used to first describe STC. A strong stimulus (in our case a sTBS) was delivered to one input (denoted S0) and 30 min later a weak stimulus (in our case a wTBS) was delivered to a second, independent input (S1). Representative sample traces are an average of four consecutive responses from typical experiments. Each experiment was from hippocampal slices of different animals, hence the n repetitions value represents both the number of slices and animals used.

### Statistical Analyses

All treatment groups were interleaved with control experiments. Data are presented as mean ± SEM (standard error of mean). In all time course plots shown, slopes of fEPSPs were normalized to the average slope fEPSP data from the 10 min baseline periods indicated unless denoted otherwise. Bar plots and cumulative distribution plots quantify the synaptic fEPSP data averaged over the 10 min periods indicated. Statistical significance between groups was determined using a one-way ANOVA followed by post hoc Dunnett’s multiple comparisons test or Tukey’s test, as appropriate. In fig. 4 the baseline and data at t = 80-90 min were compared using a paired t-test on the raw data to determine whether significant heterosynaptic potentiation was observed. All statistical tests were performed using GraphPad Prism Version 10.0.03 for macOS, GraphPad Software, Boston, Massachusetts, USA (www.graphpad.com). Levels of significance are denoted as follows: *p < 0.05, ** p<0.01 and ***p<0.001.

### Compounds

The compounds used were 3-((2-Methyl-1,3-thiazol-4-yl)ethynyl)pyridine hydrochloride (MTEP; Hello Bio Inc, USA) and 6-Amino-*N*-cyclohexyl-N,3-dimethylthiazolo[3,2-*a*]benzimidazole-2-carboxamide hydrochloride (YM 298198; YM; Hello Bio Inc, USA). Both were stored in aliquots at −30 °C, thawed prior to use and diluted into ACSF at least 20 min before their bath application.

## ACKNOWLEDGEMENTS

This research was funded by CIHR (Canadian Institutes of Health Research) Foundation Grant #154276 (to G.L.C.). GLC is also the holder of the Krembil Family Chair in Alzheimer’s Research. We thank the Dani Reiss Family Foundation (Neurodegeneration and Aging Research Program) and Robert S. (Butch) Mandel for their support.

## AUTHOR CONTRIBUTIONS

L.A.K.: conceptualization, investigation, methodology, formal analysis, visualization, writing−original draft, writing−editing; T.M.S.: conceptualization, supervision, writing−original draft, writing−review and editing; J.G.: conceptualization, supervision, writing−original draft, writing−review and editing, funding acquisition, project oversight. G.L.C.: conceptualization, supervision, writing−original draft, writing−review and editing, funding acquisition, project oversight.

